# Lateral gating mechanism and plasticity of the BAM complex in micelles and *E. coli*

**DOI:** 10.1101/2023.08.13.553113

**Authors:** Aathira Gopinath, Tobias Rath, Nina Morgner, Benesh Joseph

**Author notes:** To whom the correspondence may be addressed: Prof. Dr. Benesh Joseph, **Email:**.

## Abstract

The β-barrel assembly machinery (BAM) mediates folding and insertion of the majority of OMPs in Gram-negative bacteria. BAM is a penta-heterooligomeric complex consisting of the central β-barrel BamA and four interacting lipoproteins BamB, C, D, and E. The conformational switching of BamA between inward-open (IO) and lateral-open (LO) conformations is required for substrate recognition and folding. However, the mechanism for the lateral gating or how the structural details observed *in vitro* correspond with the cellular environment remains elusive. Here we addressed these questions by characterizing the conformational heterogeneity of BamAB, BamACDE and BamABCDE complexes in detergent micelles and or *E. coli* using pulsed dipolar electron spin resonance spectroscopy (PDS). We show that the binding of BamB does not induce any visible changes in BamA and the BamAB complex exists in the IO conformation. The BamCDE complex induces an IO to LO transition through a coordinated movement along the BamA barrel. However, the extracellular loop (L6) is unaffected by the presence of lipoproteins and exhibits a large segmental dynamics extending to the exit pore. PDS experiments with BamABCDE complex in intact *E. coli* confirmed the dynamic behavior of both the lateral gate and the L6 in the native environment. Our results demonstrate that the BamCDE complex plays a key role for the function by regulating lateral gating in BamA.

## Introduction

Gram-negative ESKAPE pathogens, which are a group of antibiotic-resistant pathogens, constitute a major threat to human health worldwide (1). They have a unique outer membrane (OM) as a selective barrier to protect against harmful external factors including antibiotics. The OM is an asymmetric bilayer consisting of phospholipids and lipopolysaccharides. In addition, it carries numerous β-barrel proteins (known as outer membrane proteins or OMPs) performing crucial biological functions. OMPs are also present in mitochondria and chloroplasts and altogether they perform a wide range of functions including transport of nutrients, signaling, motility, membrane biogenesis, protein import and secretion among others (2, 3). The proper assembly and maintenance of OMPs are crucial for bacterial survival and virulence. The folding and insertion of the majority of the OMPs are mediated by the β-barrel assembly machinery (BAM) (4-10). BAM is a penta-heterooligomeric complex composed of five components, namely BamA, BamB, BamC, BamD, and BamE (11-14). BamA, which is a 16-stranded β-barrel forms the central component (11, 15-17). The four other lipoproteins (BamB-E) interact with the polypeptide transport associated domains (POTRA 1-5 named as P1-P5) of the BamA at the periplasmic side (18-20). The unfolded OMPs (uOMPS) in complex with the chaperone interact with BAM in the periplasm (21, 22) and subsequently the uOMP is folded and inserted into the outer membrane (10, 23-25). BamA and BamD are conserved (4), whereas the precise function of the other lipoproteins remains elusive. Due to the presence of BAM complex in all Gram-negative bacteria including the ESKAPE pathogens, it is a potential target for novel antibiotics (26-30).

The crystal structure of the BamACDE complex revealed BamA in the lateral-open (LO) conformation and it was suggested that BamCDE binding induces opening of the lateral gate and exit pore in BamA (18). A subsequent structure further confirmed the LO conformation of the BamACDE complex (31). However, in the same work the BamABCDE full complex was observed in the inward-open (IO) conformation and it was suggested that BamB, but not BamCDE may regulate the lateral gating of BamA. Adding further to this ambiguity, a subsequent cryo-EM structure revealed the BamABCDE full complex in the LO conformation (32). Though the cycling between IO and LO conformations is essential for BAM function (25, 32-34), how this is achieved and regulated remains elusive.

The observations above altogether show that BamA barrel is a very dynamic structure and adopts a specific conformation in response to changes in its subunit composition and or the surroundings. Structures captured distinct states from this broad conformational space. Interestingly, when observed in the native outer membranes, BamA showed a heterogenous conformation spanning multiple states (35). A systematic observation of BamAB, BamACDE and BamABCDE complexes under identical conditions is necessary to exclude the differential effects of the environments and to elucidate the gating mechanism. The *in vitro* observations also need to be validated in the cellular environment when feasible. Here we achieved these goals by observing BamAB, BamACDE and the BamABCDE complexes in detergent micelles using pulsed electron-electron double resonance (PELDOR or DEER) spectroscopy. Further, the conformation and heterogeneity of the key structural elements in BamA were examined by observing the BamABCDE complex in intact *E. coli* using *in situ* PELDOR spectroscopy (36-38). Our results provide detailed structural and dynamic information for these complexes and show that BamCDE, but not BamB binding switches BamA into the LO conformation both *in vitro* and *in situ*.

## Results

### Observing the conformation of BAM sub/full complexes using DEER/PELDOR spectroscopy

The BamAB, BamACDE and BamABCDE complexes eluted as a monodisperse peak from the size-exclusion chromatography (SEC) column. The individual subunits were well resolved for all of them as observed from the SDS-PAGE gel (Fig. S1 A-B). We further identified these complexes using laser induced liquid bead ion desorption mass spectrometry (LILBID-MS) (39, 40). The native-MS data also revealed that BamA can form stable interactions with one to four lipoproteins in various combinations (Fig. S1C). Electron spin resonance (ESR or EPR) spectroscopy techniques constitute a versatile set of tools to study biomolecules, in particular for membrane proteins under both *in vitro* and *in situ* conditions (41-45). Conventionally, nitroxide-based spin labels are engineered at desired sites through the reaction of a thiosulfate-functionalized spin label with a cysteine residue. Here we used the methanethiosulfonate spin label (MTSL) to selectively label BamA at positions located on the extracellular-, β-barrel-and the periplasmic regions (Figs. 2-6, panels A). The distance distribution between such engineered spin pairs were determined using DEER/PELDOR spectroscopy (46, 47). We chose several positions based on the available structures such that the different conformations of BamA can be selectively observed for BamAB, BamACDE and BamABCDE complexes. A Cys-less variant of BamA was created for this purpose after substituting the two native cysteines (at positions 690 and 700) to a serine, which was shown to least affect the function (32). At the extracellular side, β16 strand (at position Q801C) was paired with loop 1 (L1, at position T434C). L1 was also connected with L6 (at position S690C) and L8 (at position G796C). The L8 was connected with L3 (at position L501C) as well. At the periplasmic side, turn 1 (T1, at position T452C) was coupled to turn 6 (T6, at position S732C). To monitor orientation of the POTRA5, position T359C was paired with turn 7 (T7, at position L780C). To further characterize the BamABCDE complex, additional spin pairs were engineered. The L1 was connected to L6 through the T434C-S657C and T434C-S700C pairs. The internal dynamics of L6 was further probed using the S690C-S700C variant. The L3 (L501C) was connected to L6 (at position S700C), L7 (at position S751C), and β16 (at position Q801C). All these variants were purified into DDM micelles and could be labeled using MTSL with high efficiency (Table S1). For observing the BamABCDE complex in *E. coli*, L1-L8 (T434C-G796C), L3-L6 (L501C-S690C, L501C-Q693C), L3-L7 (L501C-S751C), L3-L8 (L501C-G769C), and L4-L8 (D562C-G796C) variants were used. The Cys-less and or single cysteine variants were used as the control samples. All the variants supported the growth of *E. coli* cells having the native BamA expression controlled with an arabinose promoter (Fig. S2).

### BamA exhibits distinct response to lipoproteins at different positions along the barrel

The overall dynamics at the spin labeled site is encoded into the room temperature continuous wave (RT CWR) ESR spectrum. Thus, such spectra can provide information on site-specific variation of dynamics or its modulation by other interacting partners at physiological temperatures (48). We obtained RT CW ESR data using doubly labeled variants of BamAB, BamACDE and BamABCDE at a few key positions (Fig. 1 and Fig. S3). As the spectra is a superposition of signals from the two labeled sites, it gives an averaged value for the overall change in dynamics. Nevertheless, it allows for a direct comparison with the conformational changes observed from the DEER/PELDOR data (presented in the latter sections). For the lateral gate (434-801), a comparative observation of the data shows spectral narrowing for BamACDE and BamABCDE (Fig. 1). This reveals an increased dynamics as compared to BamAB, which is in line with an opening of the lateral gate in both BamACDE and BamABCDE (31, 32). For a more quantitative analysis, we extracted the corresponding correlation times (τ_c_) through a multiparameter fitting (overlaid in red dotted lines) of the spectra (49). This gave a τ_c_ of 4.5 ± 0.5 ns in BamAB, which decreased to 3.5 ± 0.4 ns for the other two complexes (Table 1).

**Figure 1.**
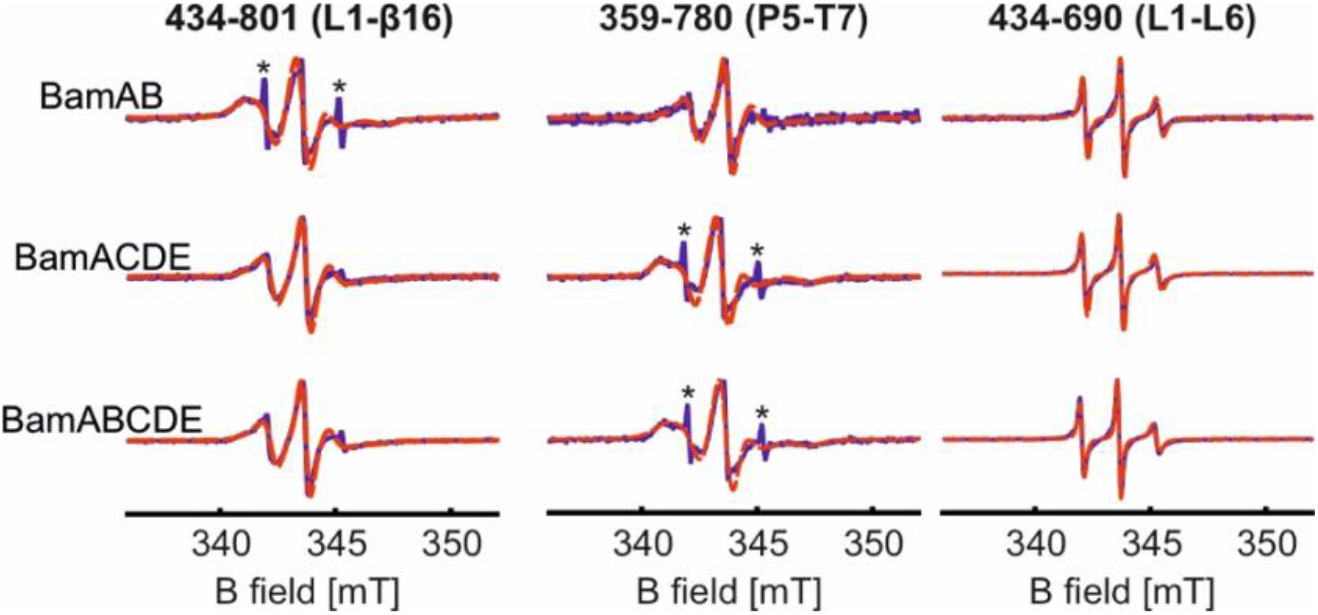
RT CW ESR spectra of the selected positions for BamAB, BamACDE and BamABCDE in DDM micelles. Simulations obtained using the EasySpin program are overlaid (in red dotted lines) and the corresponding correlation time values are given in Table 1. The asterisks indicate signal from a small fraction of free spin labels.

**Table 1.**
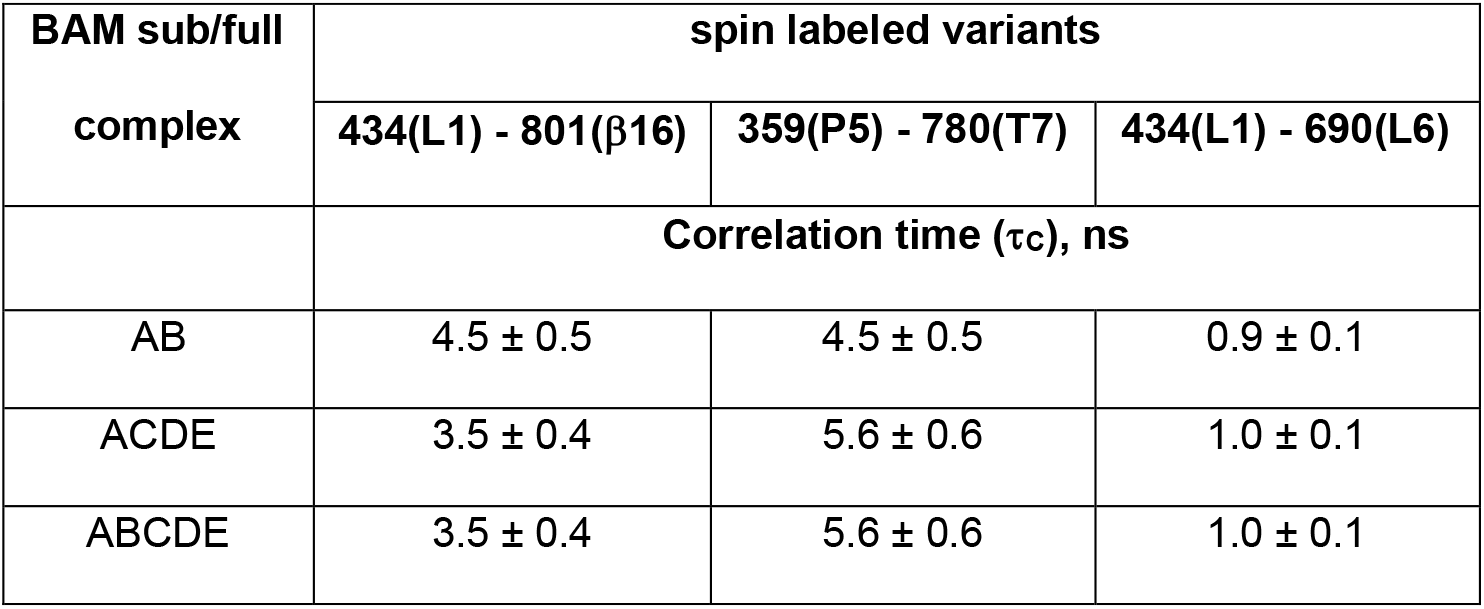
Correlation times obtained for the selected spin labeled BAM sub- and full-complex variants using RT CW ESR spectroscopy. The corresponding spectra and simulations are shown in Fig. 2.

For the P5-T7 pair (359-780) we see an opposite response where the dynamics is reduced (with increased spectral width) in the BamACDE and BamABCDE complexes (τ_c_ = 5.6 ± 0.6 ns) in relation to BamAB (τ_c_ = 4.5 ± 0.5 ns). This agrees with the P5 closed conformation for BamACDE and BamABCDE as observed in the LO structures and the dynamic changes previously observed from a solid-state NMR study (31, 32, 50). Strikingly, the L1-L6 pair (434-690) gave a very narrow spectrum revealing a significantly enhanced dynamics. Position 434 was also used to observe the lateral gate (434-801), but did not produce such a narrow component. Therefore, narrowing must be contributed by the MTSL attached to position 690 and with its extremely narrow feature it dominated the overall shape of the spectrum. Simulations gave the smallest correlation time among all the samples investigated (1.0 ± 0.1 ns). Position 690 is located (in a non-conserved insertion) on L6, which is essential for function (15, 51). A previous NMR study showed an enhanced backbone dynamics at this insertion sequence (52) and our observations reveal such behavior for the side chain as well. Overall, the dynamics and its modulation by the lipoproteins at the lateral gate and P5 suggests an IO (in BamAB) to LO (in both BamACDE and BamABCDE) transition, which very well corresponds with the conformational changes and heterogeneity characterized using PELDOR spectroscopy (Figs. 2-6).

**Figure 2.**
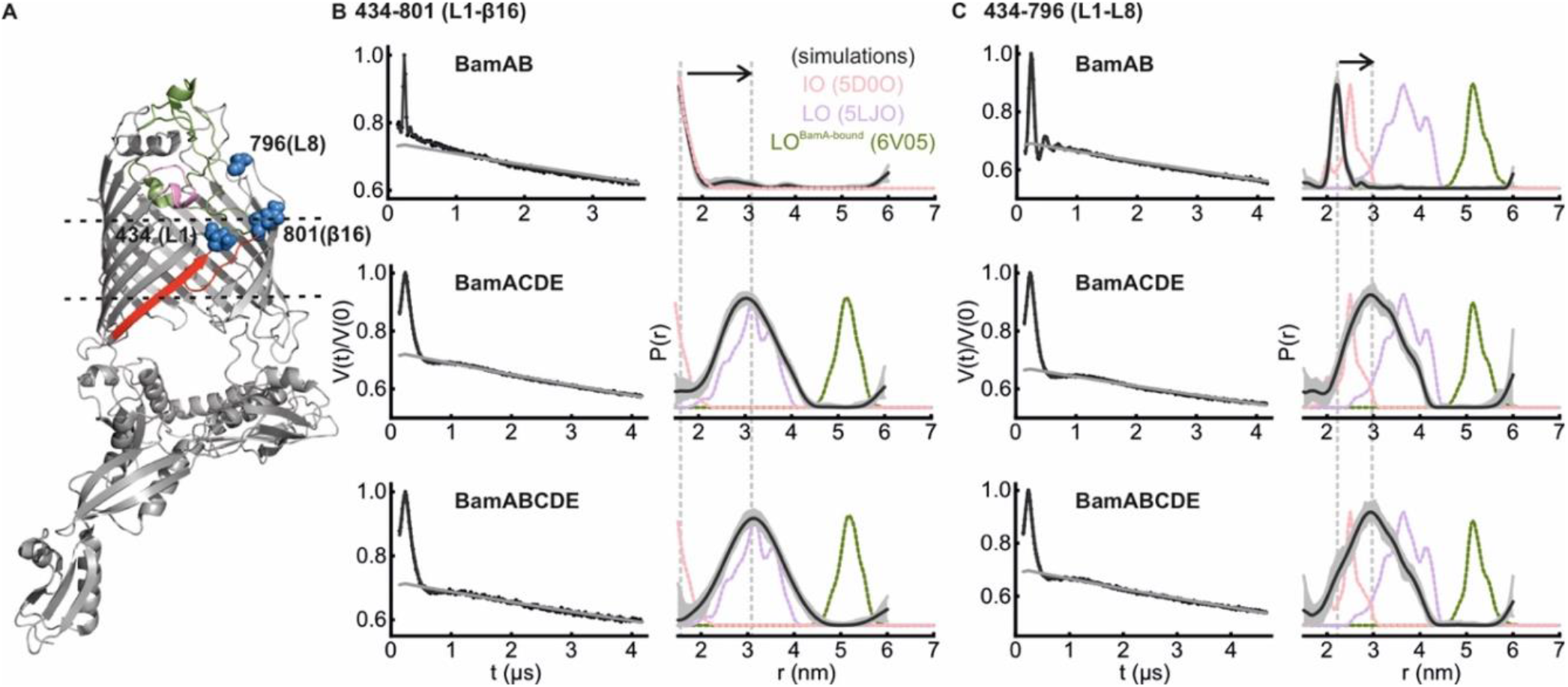
DEER/PELDOR data for the lateral gate in BamAB, BamACDE and BamABCDE in DDM micelles. (A) Structure of BamA in an inward-open conformation (PDB 5D0O). The lateral gate region (red) is closed with L3 (pink) inside the lumen and L6 (green) spanning the extracellular side. The spin labeled positions (blue) are highlighted as spheres. The lipoproteins are not shown for clarity. (B-C) Primary data overlaid with the fits obtained using the DeerLab(53) program are shown in the left panels. The obtained distance distributions with a 95% confidence interval are shown on the right. Simulations for the IO (salmon, PDB 5D0O), LO (light violet, PDB 5LJO) and LO^BamA-bound^ conformations (green, PDB 6V05) are overlaid in dotted lines.

**Figure 3.**
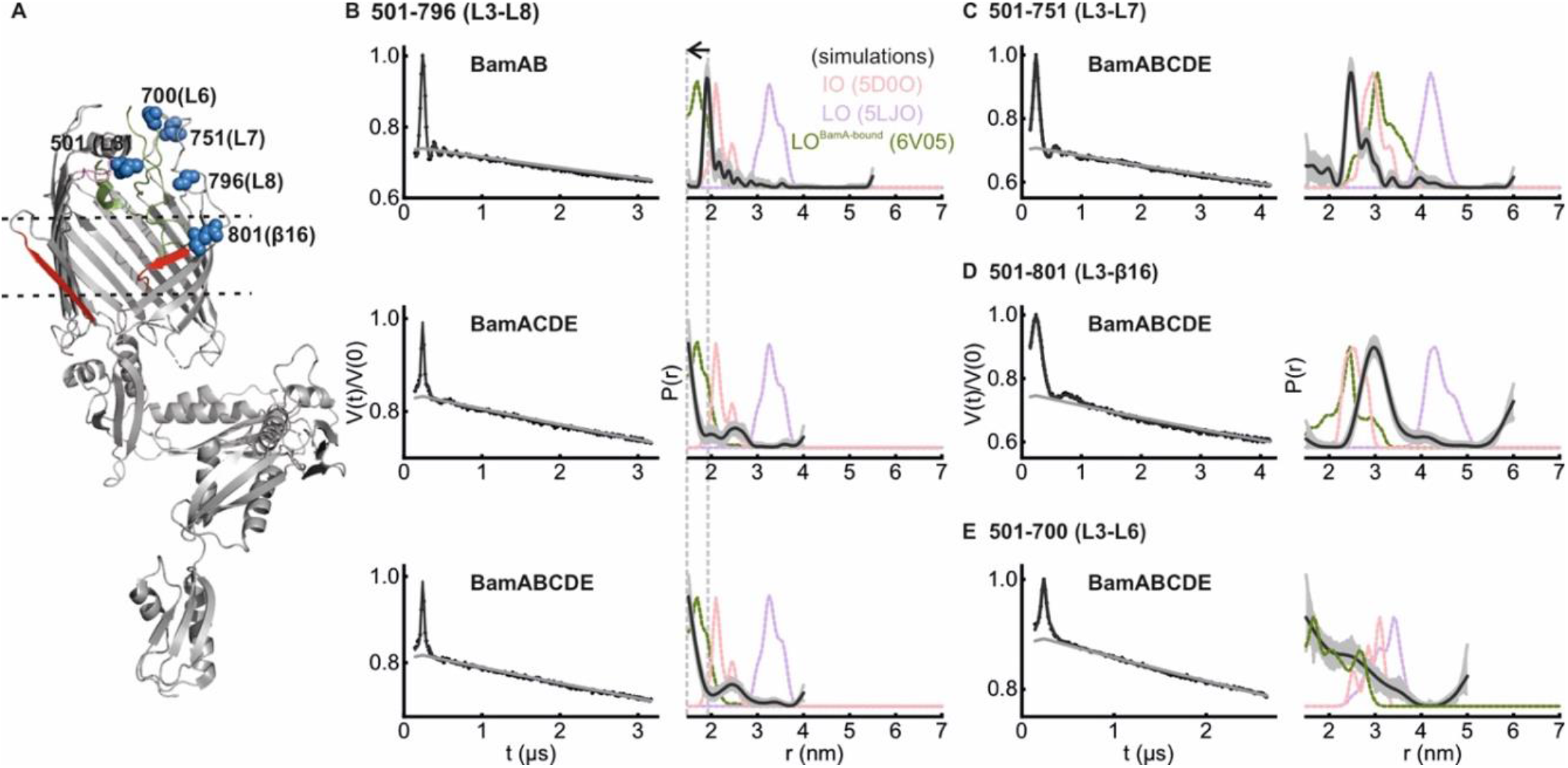
DEER/PELDOR data for the extracellular loop 3 (L3) in BamAB, BamACDE and or BamABCDE in DDM micelles. (A) Structure of BamA in the LO^BamA-bound^ conformation (PDB 6V05). The lateral gate region (red) is open to the membrane with L3 (pink) inside the lumen and L6 (green) spanning the extracellular side. The spin labelled positions (blue) are highlighted as spheres. The lipoproteins and the substrate are not shown for clarity. (B-E) Primary data overlaid with the fits obtained using the DeerLab(53) program are shown in the left panels. The obtained distance distributions with a 95% confidence interval are shown on the right. Simulations for the IO (salmon, PDB 5D0O), LO (light violet, PDB 5LJO) and LO^BamA-bound^ conformations (green, PDB 6V05) are overlaid in dotted lines.

**Figure 4.**
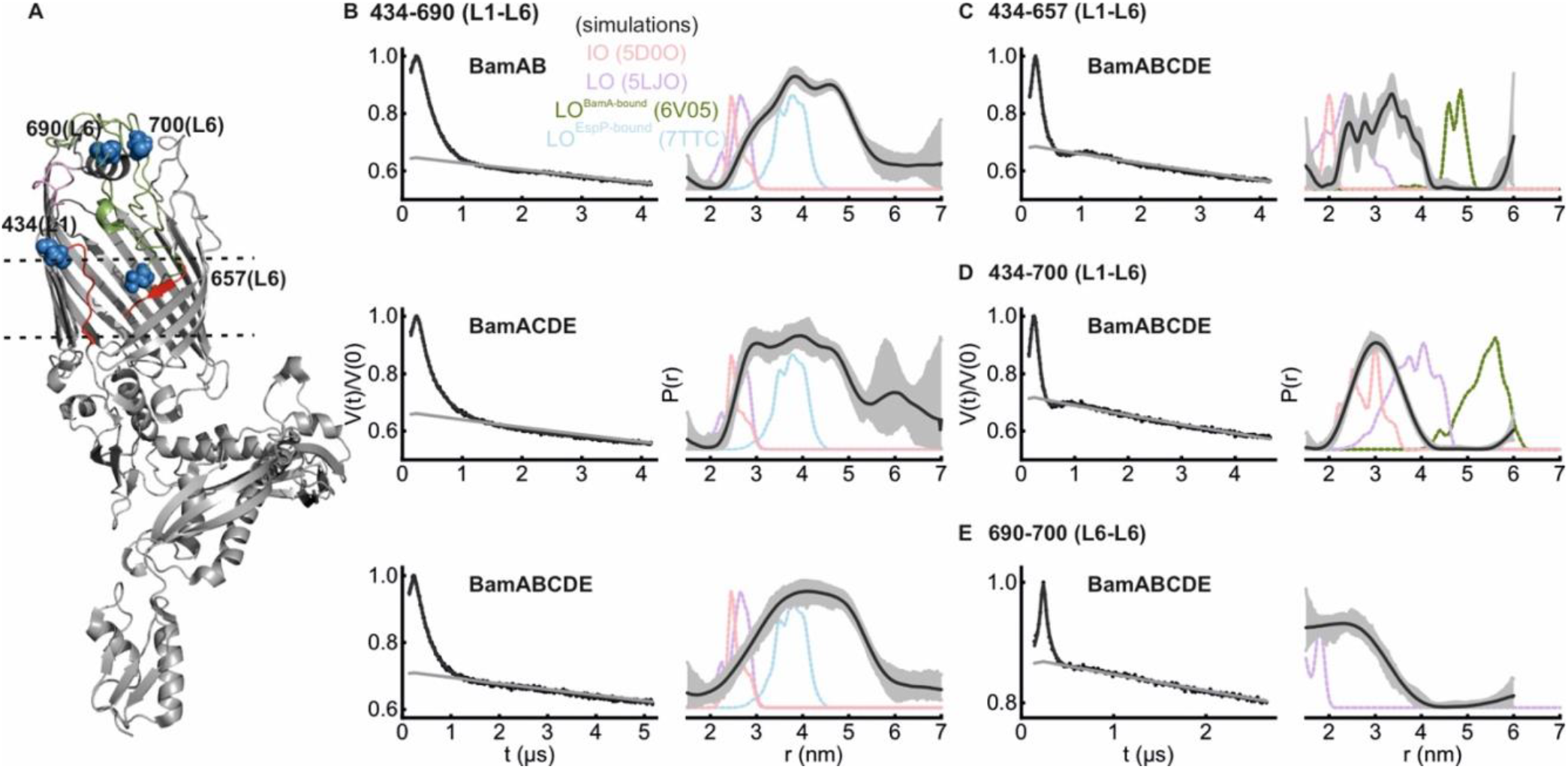
DEER/PELDOR data for the extracellular loop 6 (L6) of BamAB, BamACDE and BamABCDE in DDM micelles. (A) Structure of BamA in the LO conformation (PDB 5LJO). The lateral gate region (red) is open to the membrane with L3 (pink) facing out of the barrel and L6 (green) spanning the extracellular side. The spin labeled positions (blue) are highlighted as spheres. The lipoproteins are not shown for clarity. (B-E) Primary data overlaid with the fits obtained using the DeerLab(53) program are shown in the left panels. The obtained distance distributions with a 95% confidence interval are shown on the right. Simulations for the IO (salmon, PDB 5D0O), LO (light violet, PDB 5LJO) LO^BamA-bound^ (green, PDB 6V05) and LO^EspP-bound^ (light blue, PDB 7TTC) conformations (as possible) are overlaid in dotted lines.

**Figure 5.**
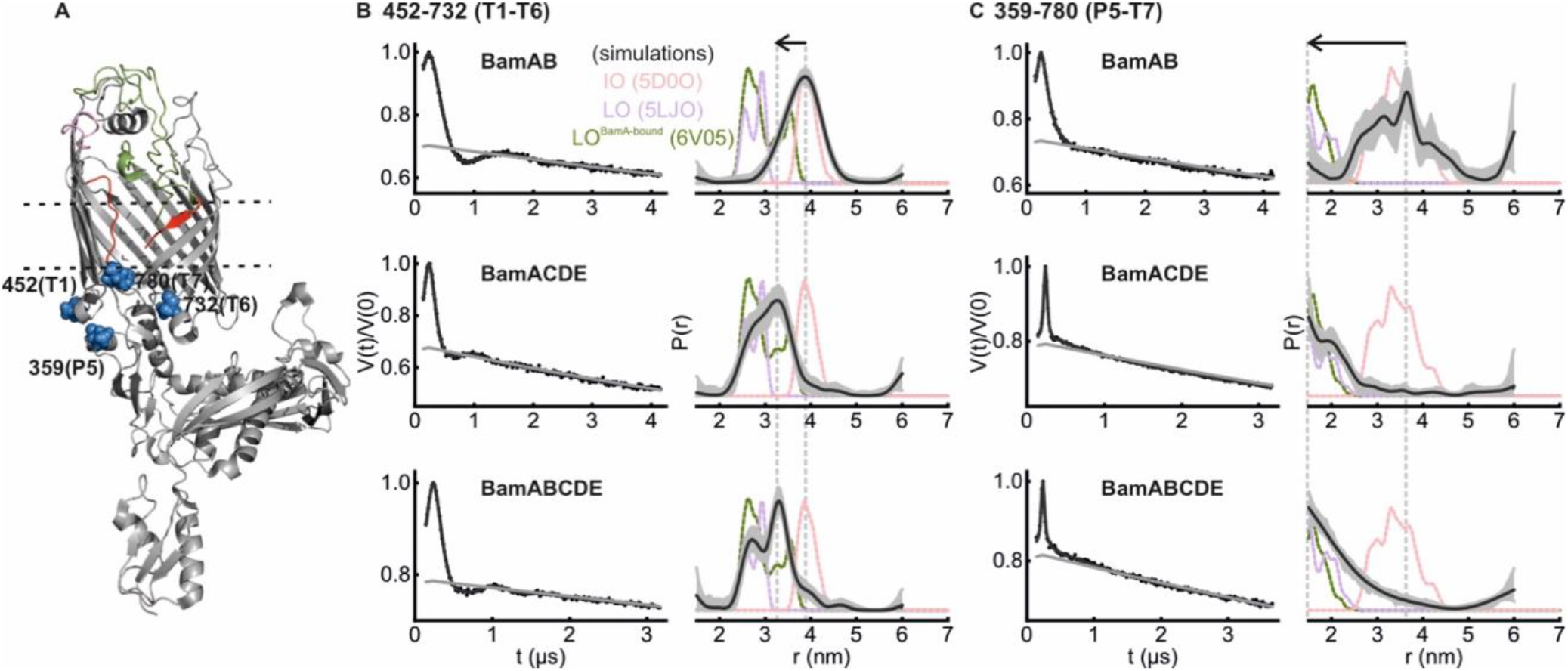
DEER/PELDOR data for the periplasmic gate (T1-T6) and between P5–T7 of BamAB, BamACDE and BamABCDE in DDM micelles. (A) Structure of BamA in the LO conformation (PDB 5LJO) as explained in Fig. 5A. (B-C) Primary data overlaid with the fits obtained using the DeerLab(53) program are shown in the left panels. The obtained distance distributions with a 95% confidence interval are shown on the right. Simulations for the IO (salmon, PDB 5D0O), LO (light violet, PDB 5LJO) and LO^BamA-bound^ conformations (green, PDB 6V05) are overlaid in dotted lines.

**Figure 6.**
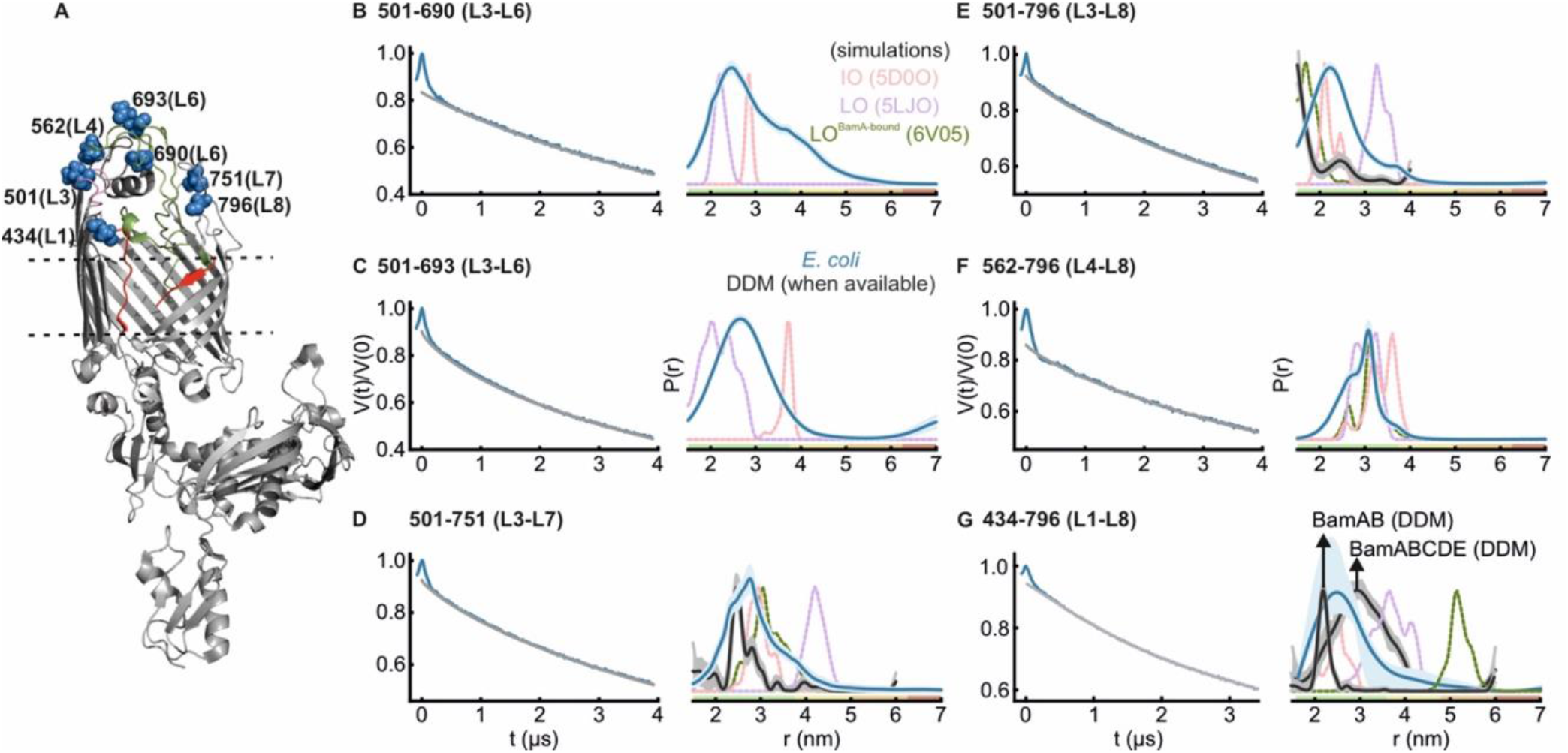
DEER/PELDOR data for the lateral gate and between the extracellular loops of BamABCDE complex in intact *E. coli*. (A) Structure of BamA in the LO conformation (PDB 5LJO) as explained in Fig. 5A. (B-F) Primary data overlaid with the background function (in grey) obtained using the DeerNet(60) program are shown in the left panels. Analysis employing Tikhonov regularization also gave a similar distance distribution (Fig. S5). (G) For the lateral gate (434-796), a polynomial background correction as determined from the corresponding single cysteine variants was employed for Tikhonov regularization using the DeerAnalysis program(61) (see Fig. S4B). The DeerNet also predicted a similar distribution except an additional small peak at longer distances, which was disregarded in view of the limited observation window of the dipolar evolution. The obtained distance distributions with an uncertainty band are shown on the right. The color code relates the reliability for different features of the probability distribution with the length of the observed dipolar evolution time. In the green zone, shape, width, and the mean distance are accurate. In the yellow zone, width and the mean, and in the orange zone, the mean distance are reliable. Distances distribution in DDM (when available) and simulations for the IO (salmon, PDB 5D0O), LO (light violet, PDB 5LJO) and LO^BamA-bound^ (green, PDB 6V05) conformations are overlaid in dotted lines. The corresponding control measurements employing single cysteine variants are shown in Figure S4. Another fully independent set of replicates produced similar results (Fig. S6-S7).

### The lateral gate is closed in BamAB and remains open in BamACDE and BamABCDE

To probe the effect of lipoproteins for the gating of BamA lateral gate, we used the 434-801 and 434-796 variants (Fig. 2A). Position 801 is located at the β16 and 796 is located five amino acids away on L8. These pairs show a large change in the interspin distances between the IO and LO conformations. For 434-801 in BamAB, the PELDOR data revealed a short interspin distance (r_max_ at 1.5 nm), which is in good agreement with simulations for the IO conformation (Fig. 2B). For BamACDE and BamABCDE, the distance significantly increased (r_max_ at 3.0 nm and 3.1 nm, respectively) in agreement with simulation for the LO conformation as well as the increased dynamics observed from the CW ESR spectra (Fig. 1). Similar results were observed for the 434-796 pair (Fig. 2C). In BamAB it gave distances close to the simulation for the IO conformation (r_max_ at 2.2 nm) and in BamACDE and BamABCDE, the mean distance increased with a broad distribution (r_max_ at 2.9 nm) similar to the LO conformation.

### L3, L7, and L8 are less flexible in detergent micelles

In the available structures, the L3 is seen to undergo large conformational changes between different conformations of BamA barrel. In the structures, the L3-L8 distances increased by more than 1 nm upon changing from the IO to the LO conformation (Figs. 2A and 4A). However, in one of the substrate-bound conformations (LO^BamA-bound^, in which the lateral gate further opened, Fig. 3A), the L3 moved more into the barrel and the L3-L8 distances decreased even below the IO conformation(23). In BamAB, the L3-L8 (501-796) PELDOR data gave a narrow distribution (r_max_ at 1.9 nm), shorter than the range predicted for the IO conformation (Fig. 3B). The narrow width of the distribution reveals a rigid orientation of both L3 and L8. Strikingly, for both BamACDE and BamABCDE in which the lateral gate is open (Fig. 2B-C), the distances further decreased (r_max_ at 1.5 nm), revealing a tightly closed conformation of L3 over the barrel lumen (similar to that observed in the LO^BamA-bound^ conformation). This is in stark contrast to the expected opening of L3 as observed in the LO structures(10, 24, 25, 31, 54). To further verify this observation, we used the L3-β16 (501-801), L3-L6 (501-700) and the L3-L7 (501-751) pairs and determined the distances in the BamABCDE complex. For L3-L7, overall, the observed distance distribution is shorter (r_max_ at 2.5 nm) than the simulation for the IO (and LO) conformation (Fig. 3C). The narrow distribution of L3-L7 distances reveal a rather rigid orientation of L7 as well. For L3-β16, the distances lie closer to the IO (and LO^BamA-bound^) conformation (r_max_ at 3.0 nm, Fig. 3D). Thus, we did not observe the open conformation of L3 in all the three variants above. Notably, the L3-L6 pair revealed a very broad distribution covering different conformations as observed in the available structures (Fig. 3E). The longer distances overlap with simulations for the IO and LO conformations, whereas a major part of the distribution is much shorter (as in the LO^BamA-bound^ structure), revealing an orientation of L3 very close to L6 over the barrel lumen. Considering the rigid orientation of L3, the broadness must be accounted by the dynamics of L6, which is also suggested from the CW ESR spectra (Fig. 1 and also see Fig. 4). Combined with the results for the lateral gate (which evidently switches between IO and LO conformations, Fig. 2), these observations reveal that BamA is a very dynamic structure and its key functional elements exhibit distinct changes in response to the presence of lipoproteins and or the nature of the surrounding environment.

### L6 is insensitive to the lipoproteins and can occupy a broad conformational space

In view of the broad L3-L6 distribution as observed above, we further investigated the conformational heterogeneity of L6 by engineering more inter- and intraloop spin pairs (Fig. 4A). The L6 forms the longest extracellular loop in BamA of *E. coli*. It covers the barrel from the extracellular side and is essential for the function of BAM complex (11). L6 contains two cysteines at positions 690 and 700, which can naturally form a disulfide bond. These cysteines were substituted in several studies, which did not affect the function (11, 32, 33). First, we investigated the effect of lipoproteins on the L6 conformation using the L1-L6 (434-690) variant. In BamAB, it gave a very broad distribution (r_max_ at 3.8 nm) spanning distances much longer than the IO and LO conformations. The lower part of the distribution corresponds to an overlay of the distances for the IO, LO and the Esp-bound (LO^EspP-bound^) conformations (24) and the remaining part reveal a continuum of conformations having L6 placed further away from L1 (Fig. 4B). Very interestingly, this distribution showed little changes when observed in BamACDE or BamABCDE, revealing that the L6 conformation is insensitive to the binding of lipoproteins. Though position 434 (L1) shows a large outward displacement during IO to LO transition, distances to position 690 (L6) do not appreciably change due to a similar relative orientation (Fig. 4B). Also, 434 (L1) distances to other positions show good agreement with the simulations (Fig. 4B-C). Thus, the broadening of L1-L6 distribution must be mostly contributed through the dynamics of L6.

In *E. coli*, residues 685-697 corresponds to a non-conserved insertion into the L6(52), which is located at the top of the barrel into the extracellular space. We designed two variants in which L6 is labeled away from the lower and upper ends of the insertion (Fig. 4C-D). One of these variants (434-657) also gave a distribution significantly broader than the (IO and LO) simulations. Position 657 is located in a region inside the barrel close to the lateral gate and forms an exit pore together with L1-L3 for the release of extracellular regions of the folding substrate. Thus, the segmental dynamics of L6 is transduced beyond the insertion sequence towards the key functional regions inside BamA. The other variant (434-700) gave distances which are comparatively less broad, revealing a reduced dynamics at position 700 (as compared to position 690). We further probed the internal dynamics of this region using wild-type (WT) BamABCDE having the native cysteines retained. A significant fraction (∼ 40%) of the molecules could be spin labeled at these positions following incubation with β-mercaptoethanol. It showed a rather broad distance distribution (Fig. 4E) and the disulfide bond might restrict this inter-residue (690-700) dynamics in the WT protein. Therefore, the removal of the native disulfide bond in L6 may account for the enhanced dynamics we observed. When the cysteines were substituted, this region had poor resolution in BamA/BAM structures(15, 16, 23, 54, 55), revealing an increased flexibility as we observed. Overall, our observations suggest that the native disulfide bond has an important role for the regulation of dynamics in L6. Though it would be informative to perform the distance measurements with WT BamA (to directly dissect the role of the disulfide bond for L6 dynamics), possible non-specific labeling of the native cysteines precluded such experiments.

### BamCDE, but not BamB binding closes periplasmic turns and POTRA 5 in BamA

We probed the response of the periplasmic turns T1 and T6 to the binding of lipoproteins using the 452-732 variant (Fig. 5A). In BamAB, the interspin distances (r_max_ at 3.9 nm) are similar to the simulation for the IO structure. However, in BamACDE and BamABCDE, the distances decreased (r_max_ at 3.3 nm) revealing the closure of these turns in line with the simulation for the LO conformation (Fig. 5B). Overall, the distribution is broader than the simulation, which reveals an increased flexibility of these turns in the LO conformation. We further probed the response of POTRA 5 using the 359-780 (P5-T7) variant. Again, in BamAB, the experimental distribution (r_max_ at 3.6 nm) is overlaid with the IO simulation. However, in BamACDE and BamABCDE, the distances were significantly decreased (r_max_ at 1.5 nm), revealing the movement of POTRA 5 towards the barrel lumen as observed in the LO conformation (Fig. 5B). This is also in line with the reduced mobility at these positions for BamACDE and BamABCDE complexes as observed from the CW ESR spectra (Fig. 1). Altogether, our observations confirm that BamCDE, but not BamB shifts BamA barrel into the LO conformation.

### Conformation of L3, L6 and the lateral gate in *E. coli*

Earlier we demonstrated that for outer membrane proteins, the structural and conformational changes can be observed in intact *E. coli* and isolated outer membranes using *in situ* spin labeling and PELDOR experiments (38, 56, 57). Solid-state NMR and single-molecule force spectroscopy techniques are complementary tools for such investigations, in particular for isolated outer membranes (58, 59). Using this approach, we showed that BamA displays an increased heterogeneity, in particular at the lateral gate when observed in the native outer membranes (35). Here we extended similar experiments for many other interloop pairs of BamABCDE full complex in intact *E. coli* cells (see Figs. S4-S8). Spin labeling is rather limited for the surface exposed cysteines in *E. coli*. The L3 was connected to L6 using the 501-690 and the 501-693 variants. Both of them gave a probability distribution significantly broader (r_max_ at 3.0 nm and 2.8 nm, respectively) than the simulations for the IO and LO conformations (Fig. 6B-C). Thus, the segmental dynamics of L6 observed in DDM micelles (Fig. 4) is preserved in the cellular environment. For the L3-L7 pair (501-751), the experimental distribution lies closer to the simulation for the IO (or LO^BamA-bound^) conformation, but is significantly broader than both the simulation and the detergent solubilized sample (Fig. 6D). In agreement, the L3-L8 (501-796) distribution is also appreciably longer and broader than the results in detergent micelles and the IO simulation (Fig. 6E). The overall distribution is more centered at the IO conformation and a small fraction of distances overlaps with the simulation for the LO conformation. For L4-L8 (at 562-796), simulations cannot differentiate between IO and LO conformations and the experimental distribution overlaps with both simulations (revealing an overall limited flexibility for both L4 and L8).

The above observation implies that the increased width for the L3-L8 (501-796, and possibly for L3-L7/501-751) distribution is attributed by an enhanced flexibility of L3. Importantly, the L1-L8 pair (434-796) reporting at the lateral gate revealed a broad distribution in *E. coli* covering the range corresponding to both IO and LO conformations, suggesting an equilibrium between closed and open conformations (Fig. 6G). Overall, the distances are significantly broader and longer than the corresponding distribution for BamAB observed in DDM micelles (Fig. 2C and overlaid in Fig. 6G). We could not verify the subunit composition of BAM for the experiments in *E. coli*. An identical expression protocol was used for the *in vitro* purification, which gave the intact BAM complex (Fig. S1A-C). Thus, BamA might interact with all of the Bam lipoproteins to form the full complex under the expression conditions. Under laboratory environment, BamA and the lipoproteins are expressed at a few thousand (1.5 – 6.0 × 10^3^) copies per cell (62). In our case, overexpression leads to a 100-fold increase in the copy number for BamA (35), which is comparable to the native expression level of some of the OMPs (such as OmpA, OmpC and OmpF). As for membrane protein overexpression in general, the cells may increase the surface area to accommodate the additional copies of BAM molecules (63). Importantly, our control experiments using single cysteine variants did not give any particular distances (Fig. S5), revealing a sufficient separation between individual BAM molecules in the native outer membrane.

## Discussion

Despite the large amount of structural data available, the exact role of the lipoproteins for the structure and dynamics of BamA remains elusive. While structures for BamA, BamACDE, BamABCDE complexes are available, BamAB or other possible subcomplex structures with BamA have not yet been determined. The heterogeneity of BamA conformation observed in various structures lead to differing ideas regarding the role of lipoproteins for BamA gating. Here we specifically addressed the role of lipoproteins on the dynamics and structure of BamA and further corroborated those observations with results obtained in intact *E. coli*.

While structures provide distinct snapshots from the conformational space, the associated changes in dynamics are obscured. The RT CW ESR spectroscopy data showed that in comparison to BamAB, the overall dynamics is increased at the lateral gate (L1-β16 / 434-801, Fig. 1) for BamACDE and BamABCDE. This is accompanied with a decreased dynamics for the P5-T7 pair (359-780). These observations agree with the IO to LO transition at the lateral gate induced by BamCDE binding. Here, the P5 moves into a closed conformation towards the barrel and the β1-β6 strands rotate outward into the LO conformation. These changes are in perfect agreement with the DEER/PELDOR results. BamCDE binding induced an opening of the lateral gate with a concomitant closure of P5 (plus T1 and T6, Figs. 2 and 5) into the barrel lumen. BamB binding alone or together with the BamACDE complex did not induce any visible changes in BamA. Thus, BamB might have an accessory role for facilitating substrate recognition, folding or interaction with the SurA chaperone in the periplasm (21).

Although the lateral gate of BamA opens upon BamCDE binding, the L3 showed an independent response. In BamAB, it gave a narrow distance distribution against L8, which is closer than the conformation observed in the IO structure. BamCDE binding further decreased the distance, revealing an even more closed conformation of L3 over the barrel and nearer to L8 as observed in the LO^BamA-bound^ structure (Fig. 3). Thus, the opening of the lateral gate (Fig. 2, as well as the closure of P5, T1-T6, Fig. 5) and the outward motion of L3 following BamCDE binding (as observed in the structures) might not be strictly correlated events. The L3 might be sensitive to the surrounding environment and the presence of the substrate, which also might explain its varying conformations between different structures. Notably, when BamA was expressed without any lipoproteins, L3 displayed a dynamic behavior spanning the range of both IO and LO conformations in the native outer membranes(35). A rather narrow interloop distance distribution against the rigid L3 reveals a limited flexibility for L7 and L8 as well (Figs. 3B-C). Such a stable orientation of L3, L7 and L8 might restrict non-specific diffusion of molecules through the barrel lumen. On the other side, the very broad distance distribution for positions on L6 revealed an extremely dynamic behavior starting at the non-conserved insertion sequence (residues 685-697) and extending towards the exit pore close to the lateral gate (Fig. 4), also in a manner fully independent to the presence of the lipoproteins (Fig. 4B, please see discussion in the next paragraph).

In *E. coli*, the L3 has a more flexible behavior, distinct from its rather rigid conformation in micelles and several structural snapshots (Fig. 6B-E). Yet, it remains nearer to the closed conformation over the barrel lumen. The enhanced segmental flexibility of L6 is preserved in *E. coli*, validating its relevance in the physiological surroundings. A previous molecular dynamics study suggested a fixed orientation of L6 in BamA (33). We did not observe BamA alone and also removed the native disulfide bond (between residues 690-700), which would increase the overall flexibility (Fig. 4E). Also, this region had poor resolution in the structures when the native cysteines were substituted (15, 16, 23, 54, 55). Thus, the native disulfide bond has a key role in regulating the overall dynamics of L6. This bond is intact in most of the available structures, however the cysteines remained non-bonded in some of them (19, 32, 64). Thus, the cysteines may also switch between bonded or non-bonded states depending on the redox state of the surrounding vicinity and thereby modulate the segmental dynamics in L6. Together with L1 and L2, L3 and L6 form a putative substrate exit pore over the BamA barrel (33). In the closed conformation, they might prevent non-specific diffusion of molecules through the barrel lumen. However, these loops move out to accommodate and release the structural elements of the substrate during protein folding (33, 65). The flexible nature of these loops might ensure that they can regulate the accessibility of the barrel lumen in a substrate-dependent manner. The well-conserved VRGF motif (residues 660-663) in L6 is located at the exit pore interface and our data (at the nearby position 657) suggest that the absence of the disulfide bond increases the flexibility of this region (Fig. 4C). This may have a direct effect on the binding and or release of substrate(s) at the lateral gate. Whether a redox transition of the native disulfide bond and the associated conformational changes have a role for function or combating challenges in the environment (such as the envelope stress due to external factors) (65) requires further investigations.

In *E. coli*, the lateral gate of BamA showed a heterogenous conformation including both open and closed conformations (Fig. 6G, 434-796). The data suggests a somewhat smaller opening as compared to the micelles. A stable opening of the lateral gate within the membrane plane might be favored through an interaction with the folding strand(s) of the substrate. This would in turn induce conformational changes at the exit pore including that of L3 (which is closed otherwise) and L6 allowing the release of the extracellular regions of the folded substrate. Although the *in situ* data qualitatively validated the conformation of L3, L6 and the lateral gate observed in micelles, several positions showed significant differences in the overall structural heterogeneity (in comparison to micelles and or the structures, Fig. 6). BAM and the surrounding lipids mutually interact leading to the modulation of their dynamics and the function of BAM (11, 34, 66-69). Thus, it would be informative to perform similar experiments in different lipid bilayers including the native outer membranes, which is unfortunately beyond the scope of the current investigation.

In summary, BamAB adopts an IO conformation and binding of BamCDE is correlated with the closure of P5, T1, T6 into the BamA lumen and opening of the lateral gate. BamD interacts directly with P5 and thereby might drive opening of the lateral gate, which is necessary for function (25, 32, 33). Thus, our observations also provide an explanation for the essentiality of BamD for the function of WT BamA, which also might interact with the substrate (70, 71). The exact role for the other lipoproteins remains to be elucidated and they might be required to facilitate and or accelerate the folding of diverse substrates in the physiological environments (5, 54, 72, 73). The possibility to observe BAM in *E. coli* and in the native outer membranes (35, 74, 75) provides a great opportunity to elucidate the dynamic basis of BAM function as well as its inhibition by novel compounds in the native asymmetric bilayers.

## Materials and Methods

### Plasmid construction and mutagenesis

The plasmid pJH114 (76) that encode the BamABCDE full-complex and pSK86(77) encoding BamAB subcomplex was provided by Marc Baldus. The plasmid pJH114 was used to create the plasmid pJH114-ΔB encoding BamACDE by deleting the *bamB* gene using the Q5 Site-directed mutagenesis kit (New England Biolabs). The *bamA* gene, which includes an N-terminal His_6_ tag and a thrombin cleavage site following the signal sequence, was custom synthesized (GeneArt, Thermo Fisher Scientific) and inserted into the pCDFDuet-1 vector to produce the *bamA*/pCDFDuet-1 plasmid (35). The *bamB* from pSK86 was cloned into pETDuet-1 vector to create the *bamB*/pETDuet-1 plasmid. The native cysteines (C690 and C700) of *bamA* were substituted to a serine to create the Cys-less variant. Cysteines were further introduced in *bamA* at the desired positions using the Q5 Site-directed mutagenesis kit.

### Protein expression

The BamABCDE, BamACDE and BamAB was expressed and purified using protocol adapted from previous publications (64, 77). For BamABCDE and BamACDE expression, the plasmids pJH114 and pJH114-ΔB was transformed to BL21(DE3) competent cells. Cells were grown in Lysogeny broth (LB) media containing 50 μg/mL ampicillin at 37 °C until an OD_600_ of 0.6-0.8. The culture was then diluted 1:100 times to 2x YT media (16 g/L tryptone, 10 g/L yeast extract and 5 g/L NaCl) and grown at 37 °C until an OD_600_ of approximately 0.7 is reached. Protein expression was induced with 0.4 mM IPTG and grown further for 1.5 hours. For BamAB expression, the plasmids *bamA*/pCDFDuet-1 and *bamB*/pETDuet-1 were co-transformed to BL21(DE3) competent cells. Expression was carried out in a similar manner in presence of 50 μg/mL streptomycin and 50 μg/mL ampicillin.

### Protein purification and spin labeling

Cell cultures obtained were spun down at 8000x*g* for 10 min and resuspended in 20 mL lysis buffer (20 mM Tris-HCl pH 8.0, 100 μg/mL lysozyme, 1 mM PMSF and 1 μg/mL DNaseI) per gram of cells. Cells were lysed using sonication and then pelleted down at 10,000x*g* for 20 minutes. 0.5 % N-Laurylsarcosine sodium salt was added to the supernatant to solubilize the inner membrane and stirred at room temperature for 10 minutes. The solution was then ultracentrifuged at 200,000x*g* for 1.5 hours. The resulting outer membrane pellet was resuspended in the B20 buffer (20 mM Tris-HCl pH 8.0, 300 mM NaCl, 20 mM imidazole, 1% DDM and 1 mM PMSF, 100 mL / g cells) for 1 hour. The suspension was then ultracentrifuged at 200,000x*g* for 30 minutes. The supernatant was incubated with 2 mL Ni-Sepharose High Performance slurry (GE Healthcare) for 1 hour. The mixture was then loaded onto a PD-10 Empty column (GE Healthcare) and washed with 2.5 column volumes B20 buffer containing 5 mM β-mercaptoethanol followed by 5-7 column volumes of B30 buffer (20 mM Tris-HCl pH 8.0, 150 mM NaCl, 30 mM imidazole, 0.1% DDM, 5 mM β-mercaptoethanol). The protein was eluted using 5 column volumes of B200 buffer (20 mM Tris-HCl pH 8.0, 150 mM NaCl, 200 mM imidazole, 0.1% DDM, 5 mM β-mercaptoethanol). The eluted protein was buffer exchanged to 20 mM Tris-HCl pH 8.0, 150 mM NaCl, 0.1% DDM using PD-10 Desalting column (GE Healthcare). The desalted protein was immediately mixed with 40-fold excess of the spin label, 1-oxyl2,2,5,5-tetramethyl-3-pyrroline-3-methyl methanethiosulfonate (MTSL, Toronto Research Chemicals) and kept stirring at room temperature for 30 minutes. The labeled protein was then concentrated in Vivaspin 6 concentrators (MWCO 50,000 Da for BamAB and BamACDE, and MWCO 100,000 Da for ABCDE) and subsequently applied to a Superdex 200 Increase 10/300 GL column (GE Healthcare). The eluted protein fractions were further concentrated to 12-30 μM before sample preparation.

### Protein expression and spin labeling in *E. coli*

For *in-situ* measurements, the plasmid pJH114 encoding the genes for BamABCDE was transformed into *E. coli* BL21(DE3) cells. A preculture of 20 mL LB media containing 50 μg/mL ampicillin was prepared and grown at 37 °C until an OD_600_ of 0.6. 1 mL of the preculture was inoculated to 100 mL of 2x YT media and grown further to an OD_600_ of approximately 0.7. Cells were then induced with 0.4 mM IPTG and grown for 1.5 hours. The OD_600_ of the culture was measured. An appropriate amount of culture was taken out, spun down at 7000x*g* for 10 minutes and resuspended in 40 mL cold MOPS-NaCl buffer (50 mM MOPS pH 7.5, 60 mM NaCl) to a final OD_600_ of 0.5. Cells were labelled by incubating with 10 μM MTSL for 15 minutes at room temperature with mixing. Cell suspension was then pelleted down at 7000x*g* for 10 minutes. To remove the excess MTSL, cells were washed by pelleting and resuspending in 1.5 mL buffer twice. For CW ESR measurement, the final cell pellet was suspended in approximately 40 μL buffer. For PELDOR measurements, the cells were made to a final volume of approximately 30 μL using buffer and 15% *d*_*8*_-glycerol.

### *In vivo* complementation assay

The effect of cysteine substitutions were checked using *in-vivo* complementation in JCM166 (4) cells provided by Sebastian Hiller. The plasmid *bamA*/pCDFDuet-1 containing the substitutions were transformed to JCM166 cells and plated on LB agar containing 50 μg/mL spectinomycin and 0.05% arabinose. A single colony from the plate was inoculated to 5 mL LB media containing 50 μg/mL spectinomycin and 0.05% arabinose and grown at 37 °C until the OD_600_ reached approximately 1.5. Cell culture was then washed twice to remove the arabinose by pelleting down at 4000x*g* for 10 min and resuspending in LB media. Cells were resuspended in LB to a final OD_600_ of 1.0 and serial dilutions of 10, 100, 1000, 10000, 100000 folds were prepared. 2 μL of each dilution were then spotted onto LB agar plates containing 50 μg/mL spectinomycin with or without 0.05% arabinose (35, 78).

### LILBID-MS

For mass spectrometric analysis by LILBID-MS (40) the samples were buffer exchanged into 20 mM Tris HCl, 50 mM NaCl, 0.1% DDM, pH 8.0. For each measurement 4 μL of the 10 μM sample were directly loaded into a piezo driven droplet generator (MD-K-130, Microdrop Technologies GmbH, Germany). Droplets of around 50 μm diameter are produced by this generator with a frequency of 10 Hz at 100 mbar. These droplets are transferred into vacuum and irradiated by an IR laser, leading to an explosive expansion of the droplet and a release of the solvated ions. The IR laser runs at a wavelength of 2.8 μM and was set to a maximum energy output of 23 mJ per pulse with a pulse length of 6 ns. The released ions were accelerated by a Wiley-McLaren type ion optic for analysis by a home build time-of-flight setup. The voltage in the ion source was set to -4.0 kV between the first (repeller) and the second lenses. The third plate was grounded. Between 5-20 μs (delayed extraction time) after the irradiation the repeller was pulsed to -6.6 kV for 370 μs. The reflectron was set to -7.2 kV. The detector is a Daly-type, optimized for high m/z. Processing of spectra was done by using Massign, a software based on Labview(39).

### RT CW ESR spectroscopy

A Bruker EMXnano benchtop spectrometer operating at the X-band frequency was used to conduct continuous wave ESR measurements at room temperature. A 20-40 μL sample was used in a micropipette (BRAND, Germany, with a diameter of 0.68, 0.86 or 1.2 mm). The CW ESR spectra were acquired with 100 kHz modulation frequency, 0.6-2 mW microwave power, 0.15 mT modulation amplitude, and 18 mT sweep width. *In situ* samples were subjected to signal averaging over 40 scans, while protein samples were averaged over 50 to 150 scans.

### Pulsed ESR spectroscopy and data analysis

Pulsed ESR experiments were performed on a Bruker Elexsys E580 Q-Band Pulsed ESR spectrometer with SpinJet AWG equipped with an arbitrary waveform generator (AWG), a 50 W solid state amplifier, a continuous-flow helium cryostat, and a temperature control system (Oxford Instruments). A 15-20 μL sample containing 15% *d*_*8*_-glycerol was transferred into a 1.6 mm outer diameter quartz EPR tube (Suprasil, Wilmad-LabGlass) and snap-frozen in liquid nitrogen. Measurements were performed at 50 K using a dead-time free four-pulse sequence and a 16-step phase cycling (x[x][x_p_]x) (79). A 38 ns Gaussian pulse set to the maximum of the echo-detected field swept spectrum was used as the pump pulse. The 48 ns Gaussian observer pulses were set at 80 MHz lower than the pump pulse. The deuterium modulations were averaged by increasing the first interpulse delay by 16 ns for 8 steps. The dipolar evolution time window was adjusted based on the observed phase memory time *T*_*M*,_ which was determined using 48 ns π/2– τ– π Gaussian pulses and a two-step phase cycling, while τ was increased in 4 ns steps.

Data analysis was performed using the DeerAnalysis 2022, DeerNet or the DeerLab program as specified (53, 60, 61). For DeerLab, the primary data was fitted with a non-parametric distribution and a homogenous background using Tikhonov regularization (TR) and the uncertainty was estimated using bootstrapping. Data analysis employing TR was performed as implemented in the MATLAB-based DeerAnalysis 2022 package. The background function arising from the intermolecular interactions was removed from the primary data V(t)/V(0) to obtain the form factor F(t)/F(0). The resulting form factor was fitted with a model free TR to obtain the distance distribution. Error estimation of the probability distribution was determined using the validation procedure wherein the background time window and or the dimensionality of the spin distribution was gradually changed (see Table S2). Data analysis was also performed with a user independent approach employing deep neural network (DeerNet Spinach SVN Rev 5662) as implemented in DeerAnalysis package. Distance distributions were simulated employing a rotamer library approach using the MATLAB based MMM2021.1 software package(80).

BAM structures in the inward open (PDB 5D0O), lateral open (PDB 5LJO) and substrate-bound (PDB 6V05 and 7TTC) conformations were used for the simulations.

## Acknowledgments

This work was financially supported from the Deutsche Forschungsgemeinschaft via the Emmy Noether program (JO 1428/1–1), SFB 1507 – “Membrane-associated Protein Assemblies, Machineries, and Supercomplexes”, and a large equipment fund (438280639) to B.J. N.M acknowledges support from Deutsche Forschungsgemeinschaft (426191805). A.G. and B.J. thank Marc Baldus for providing pJH114 and pSK86 plasmids and initial expression/purification protocols and Sebastian Hiller for providing *E. coli* JCM166 cells.

## Author contributions

A.G. performed all the biochemical and ESR spectroscopy experiments. T.R. and N.M. performed and analyzed the LILBID-MS experiments. A.G. and B.J. designed the experiments, analyzed the data and wrote the paper with inputs from T.R. and N.M. B.J. conceived the idea, acquired funding, supervised and administered the project.

