## Supplemental Information for "Lateral gating mechanism and plasticity of the BAM complex in micelles and *E. coli*"

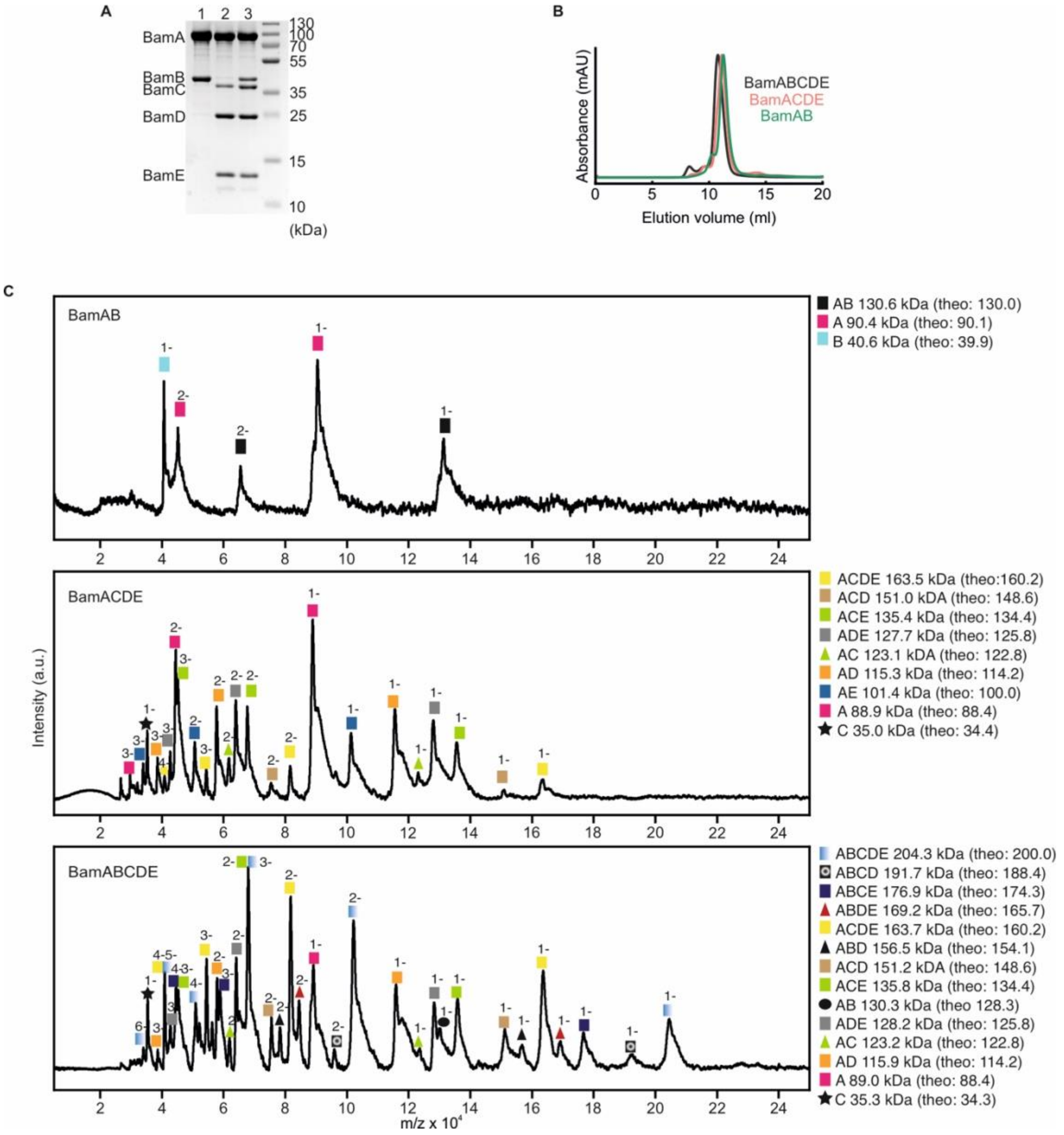


**Figure S1. Characterization of full- and sub-complexes of BAM**. (A) SDS-PAGE of the purified protein complexes, BamAB (lane 1), BamACDE (lane 2) and BamABCDE (lane 3) in detergent micelles. (B) Size-exclusion chromatogram of BamABCDE (dark grey), BamACDE (salmon) and BamAB (green). (C) LILBID mass spectrometry of BAM complexes. BamAB in the upper panel, BamACDE in middle panel and BamABCDE in the lower panel. Peaks were assigned based on the predicted molecular mass (indicated as theo:) of the complexes.

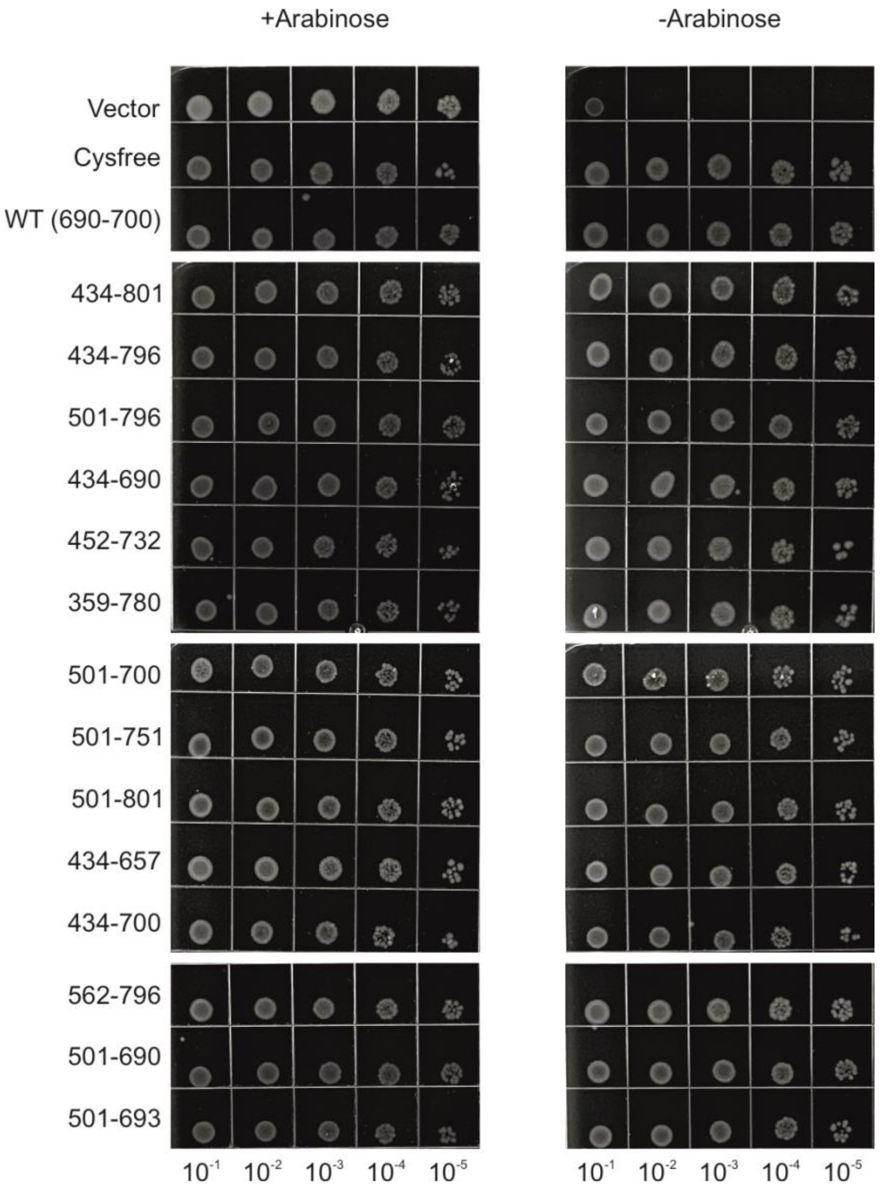


**Figure S2. Colony growth assay for BamA cysteine variants in *E. coli* JCM166 cells**. The cysteine variants expressed from the corresponding plasmids showed similar growth in the presence (on the left) and absence of arabinose (on the right). Cells were diluted for each variant as indicated. Cells transformed with the empty pCDFDuet-1 vector as a negative control did not grow in the absence of arabinose (see the Methods section for details).

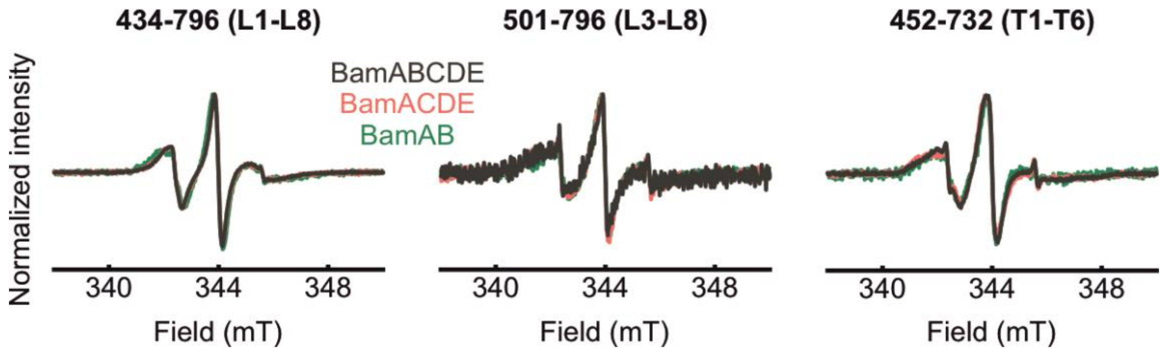


**Figure S3. Room temperature CW ESR spectra of the spin labelled double cysteine variants in DDM micelles**. The different oligomeric states BamAB (green), BamACDE (salmon) and BamABCDE (dark grey) for each variant is overlaid. Spectra are normalized for a direct comparison of the shape and spectra for the other variants are shown in Fig. 1.

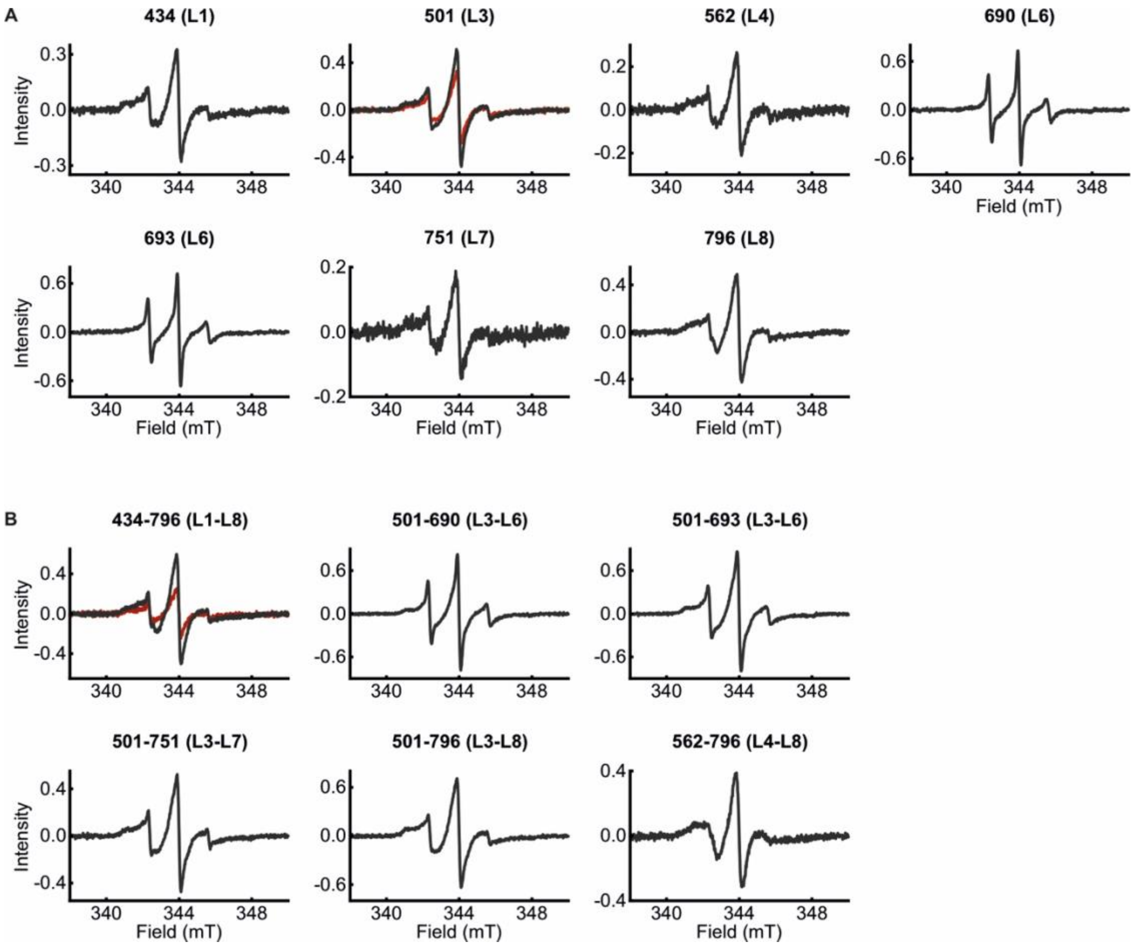


**Figure S4. Room temperature CW ESR spectra of the spin labeled single cysteine (A) and double cysteine (B) variants of the BamABCDE complex in *E. coli***. The labeled positions and the corresponding loops are indicated. The single cysteine variants were used as the control(s) for DEER/PELDOR experiments with double cysteine variants (see Figs. S5-S8). The background labeling is shown by the Cys-less BamABCDE (overlaid in red) when available. Spectra are normalized to the same cell density ($OD_{600}$) value for a quantitative comparison. The overall spin concentration obtained was in the range between 30-100 μM (see the methods section on expression and spin labeling in *E. coli*).

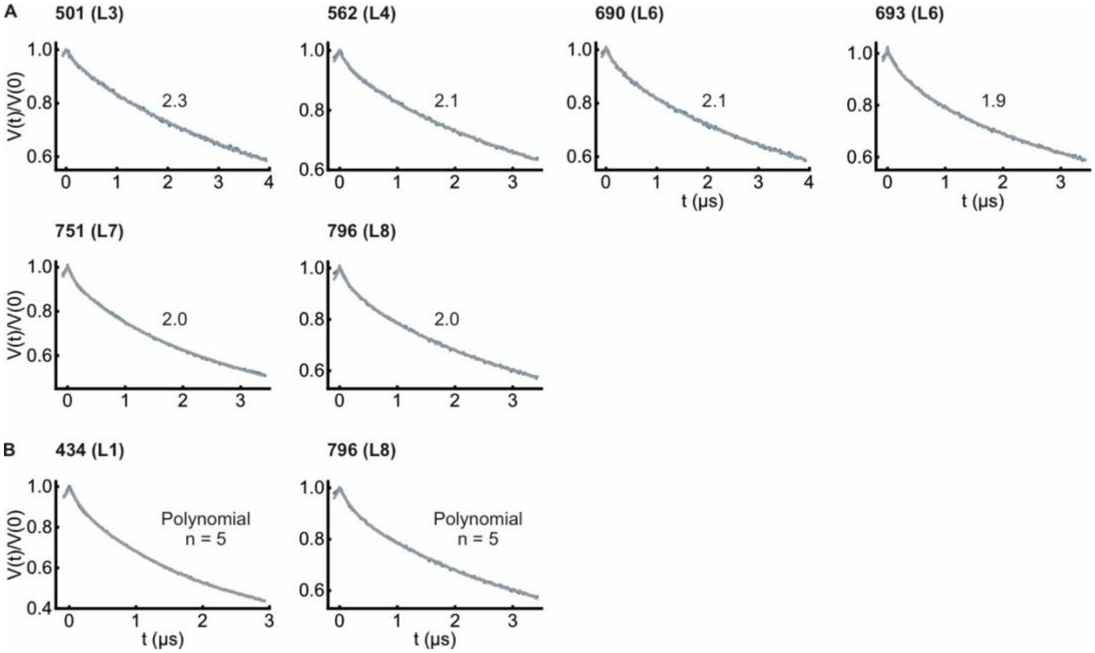


**Figure S5. DEER/PELDOR measurement of spin labeled single cysteine variants of the BamABCDE complex in *E. coli***. (A) The primary data (blue) and the fit (grey) obtained using a stretched exponential decay corresponding to the dimensionality for spin distribution (*d*) as indicated. (B) The primary data (blue) and the fit (grey) obtained by fitting the decay to a polynomial function (n=5, which together corresponded to a stretched exponential decay with *d* = 2.0-2.5).

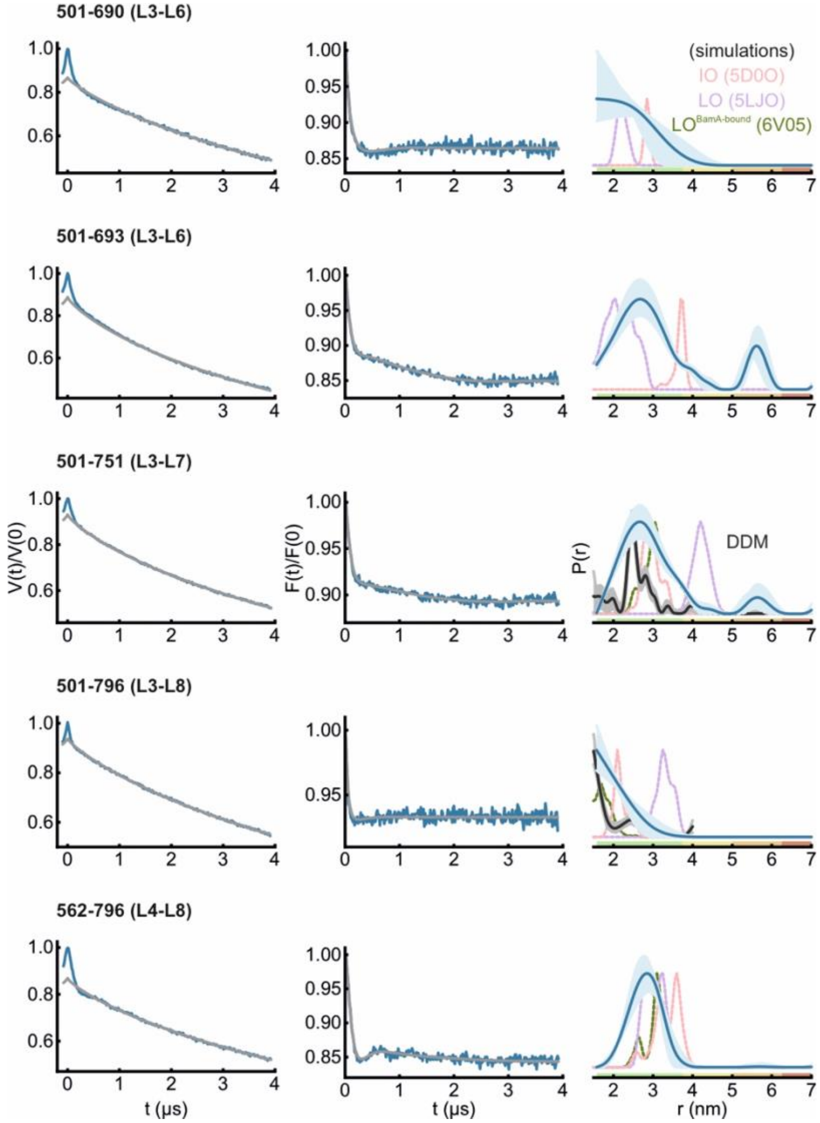


**Figure S6. DEER/PELDOR data for BamABCDE in *E. coli* analyzed using Tikhonov regularization**. Corresponding analysis using the DeerNet program is shown in Fig. 6. Primary data (blue) with the intermolecular contribution (grey) obtained using the DeerAnalysis (1) program are shown in the left panels. The middle panels indicate the form factor obtained after intermolecular background correction. The distance distribution obtained using Tikhonov regularization are shown on the right. Simulations for the IO (salmon, PDB 5D0O), LO (light violet, PDB 5LJO) and LO$^{BamA-bound}$ conformation (green, PDB 6V05) are overlaid in dotted lines. The corresponding distance distribution (when available) in detergent micelles (dark grey, from Fig. 3B and 3C) are overlaid. The error bounds show the variation in the probability amplitude due to uncertainties in background function as described in Table S2. The color coding for the distance distribution is as explained in Fig. 6.

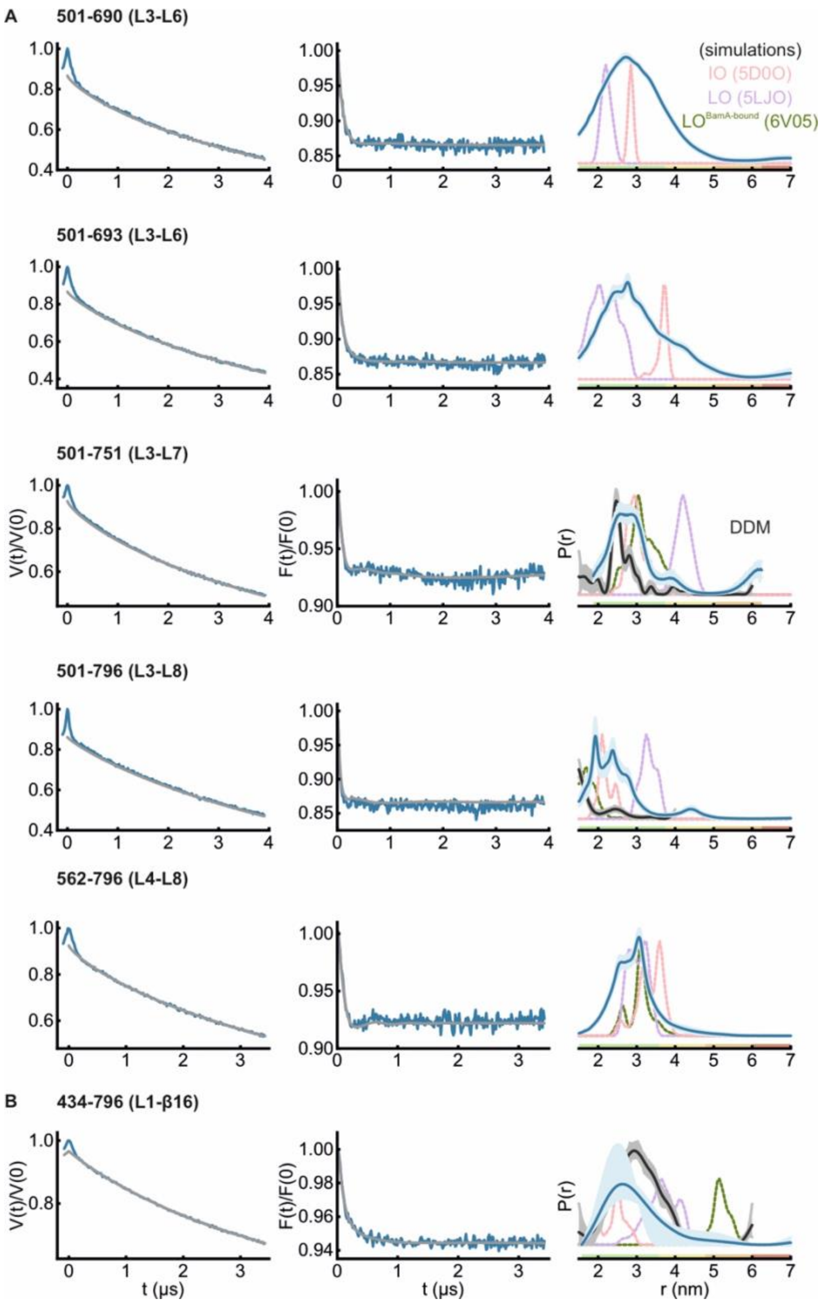


**Figure S7. DEER/PELDOR data for the replicate samples of BamABCDE in *E. coli*.** A similar data set is shown in Fig. 6 (and Fig. S6). Primary data (blue) with the intermolecular contribution (grey) obtained from the DeerNet program (A, (2)) or from the Tikhonov regularization using the DeerAnalysis program (B) as described in Fig. 6G are shown in the left panels. The middle panels show the form factors and the obtained distance distributions are shown on the right. The error bounds show the variation in distances due to uncertainties in the intermolecular function. The color code for the probability distribution is as explained in Fig. 6. Corresponding distances in micelles (DDM, when available) and simulations for the IO (salmon, PDB 5D0O), LO (light violet, PDB 5LJO) and $LO^{BamA\text{-}bound}$ conformation (green, PDB 6V05) are overlaid in dotted lines.

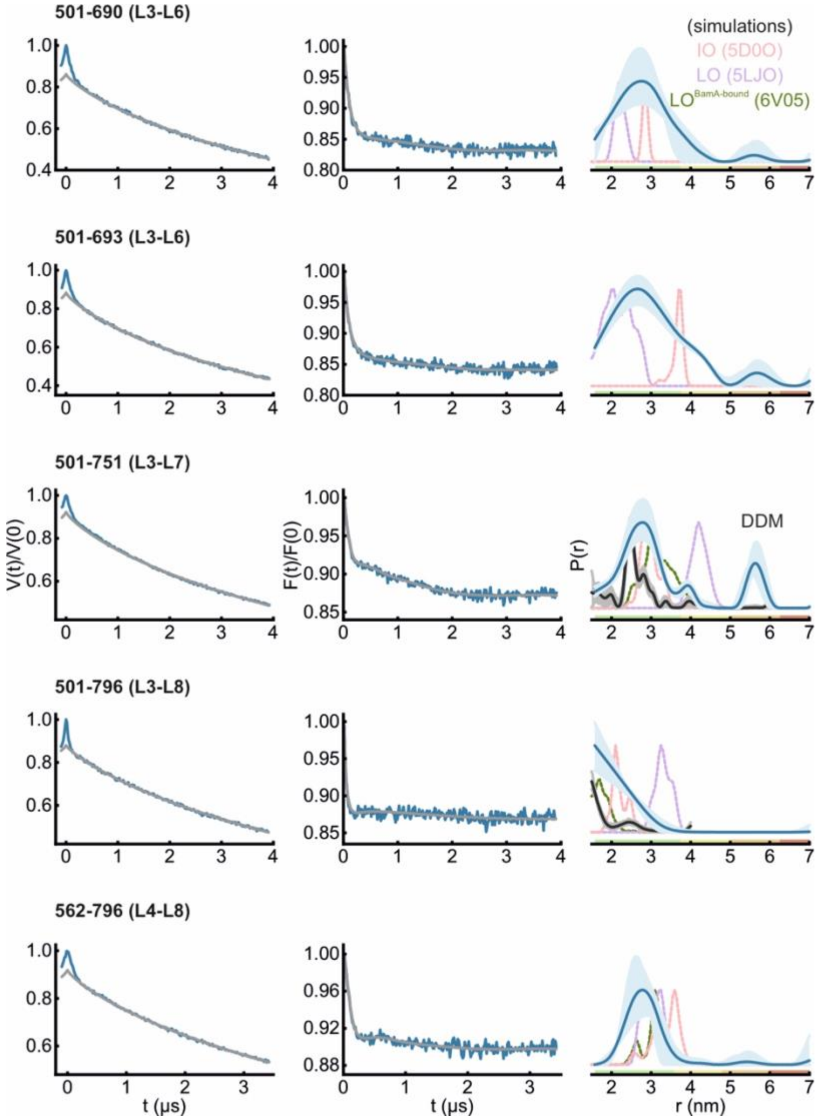


**Figure S8. DEER/PELDOR data of replicate samples of BamABCDE in *E. coli* analyzed using Tikhonov regularization.** The DeerNet analysis of the data are shown in Fig. S7. Primary data (blue) with the intermolecular contribution (grey) obtained using the DeerAnalysis program are shown in the left panels. The middle panels show the corresponding form factors. The distance distributions are shown on the right. The error bounds show the variation for the probability amplitudes due to uncertainties in background function as described in Table S2. The color code for the probability distribution is as explained in Fig. 6. Corresponding distances in micelles (DDM, when available) and simulations for the IO (salmon, PDB 5D0O), LO (light violet, PDB 5LJO) and $LO^{BamA\text{-}bound}$ conformation (green, PDB 6V05) are overlaid in dotted lines.

**Table S1.** Spin labelling efficiency for the cysteine variants of BamAB, BamACDE and BamABCDE in DDM micelles.

| Variant | Bam sub/full-complex | Protein (μM) | Spin (μM) | Labelling efficiency (%) |
|---|---|---|---|---|
| 434-801 | AB | 27 ± 5 | 51 ± 10 | 95 ± 19 |
| | ACDE | 36 ± 7 | 69 ± 14 | 97 ± 19 |
| | ABCDE | 23 ± 5 | 43 ± 9 | 94 ± 19 |
| 434-796 | AB | 25 ± 5 | 54 ± 11 | 107 ± 21 |
| | ACDE | 22 ± 4 | 45 ± 9 | 100 ± 20 |
| | ABCDE | 24 ± 5 | 51 ± 10 | 106 ± 21 |
| 501-796 | AB | 12 ± 2 | 24 ± 5 | 97 ± 19 |
| | ACDE | 24 ± 5 | 50 ± 10 | 105 ± 21 |
| | ABCDE | 17 ± 3 | 35 ± 7 | 102 ± 21 |
| 434-690 | AB | 19 ± 4 | 46 ± 9 | 124 ± 25 |
| | ACDE | 21 ± 4 | 45 ± 9 | 109 ± 22 |
| | ABCDE | 24 ± 5 | 52 ± 10 | 108 ± 22 |
| 452-732 | AB | 18 ± 4 | 41 ± 8 | 112 ± 22 |
| | ACDE | 33 ± 7 | 69 ± 14 | 103 ± 20 |
| | ABCDE | 36 ± 7 | 77 ± 15 | 108 ± 22 |
| 359-780 | AB | 16 ± 3 | 37 ± 7 | 114 ± 23 |
| | ACDE | 29 ± 5 | 52 ± 10 | 89 ± 18 |
| | ABCDE | 27 ± 5 | 63 ± 13 | 116 ± 23 |
| 501-751 | ABCDE | 20 ± 4 | 49 ± 10 | 120 ± 24 |
| 501-801 | ABCDE | 21 ± 4 | 41 ± 8 | 96.0 ± 19 |
| 501-700 | ABCDE | 21 ± 4 | 16 ± 3 | 37 ± 7 |
| 434-657 | ABCDE | 22 ± 4 | 47 ± 9 | 107 ± 21 |
| 434-700 | ABCDE | 29 ± 5 | 63 ± 13 | 109 ± 22 |
| 690-700 | ABCDE | 25 ± 5 | 18 ± 4 | 37 ± 6 |

**Table S2**. Error estimation of probability amplitudes for the samples analyzed using the DeerAnalysis program employing Tikhonov regularization. The variation of the time window and or the dimensionality for spin distribution (*d*, as determined using single cysteine variants, Fig. S5) and the regularization parameter ($\alpha$) employed for each sample is indicated.

| Variant | Figure | Error estimation | | | | | Regularization parameter (α) |
|---|---|---|---|---|---|---|---|
| | | Dimensionality (*d*) | | Starting time window | | | |
| | | range | steps | $t_{max}$ (μs) | Range (ns) | steps | |
| 501-690 | S6 | 2.0-2.5 | 6 | 4.0 | 784-2352 | 11 | 2510 |
| | S8 | 2.0-2.5 | 6 | 4.0 | 784-2336 | 11 | 501 |
| 501-693 | S6 | 2.4-2.6 | 3 | 4.0 | 784-2336 | 11 | 251 |
| | S8 | 2.4-2.6 | 3 | 4.0 | 784-2352 | 11 | 794 |
| 501-751 | S6 | 2.4-2.6 | 3 | 4.0 | 784-2352 | 11 | 501 |
| | S8 | 2.4-2.6 | 3 | 4.0 | 784-2352 | 11 | 200 |
| 501-796 | S6 | 2.2-2.5 | 4 | 4.0 | 784-2336 | 11 | 3980 |
| | S8 | 2.2-2.5 | 4 | 4.0 | 784-2336 | 11 | 2510 |
| 562-796 | S6 | 2.0-2.5 | 6 | 4.0 | 784-2352 | 11 | 501 |
| | S8 | 2.0-2.5 | 6 | 3.5 | 688-2048 | 11 | 316 |
| 434-796 | S6 | - | - | 3.5 | 500-1000 | 11 | 501 |
| | S8 | - | - | 3.5 | 500-1000 | 11 | 501 |
